# Rational Selection of PCR Primer/Probe Design Sites for SARS-CoV-2

**DOI:** 10.1101/2021.04.04.438420

**Authors:** Divya RSJB Rana, Nischal Pokhrel, Santosh Dulal

## Abstract

Various reports of decreased analytical sensitivities of real-time PCR-based detection of Coronavirus Disease 2019 (COVID-19) have been associated with occurrence of mutations in the target area of primer/probe binding. Knowledge about propensities of different genes to undergo mutation can inform researchers to select optimal genes to target for the qPCR design. We analyzed supplementary data from over 45 thousand SARS-CoV-2 genomes provided by Mercatelli et al to calculate the unique and prevalent mutations in different genes of SARS-CoV-2. We found that non-structural proteins in the ORF1ab region were more conserved compared to structural genes. Further factors which need to be relied upon for proper selection of genes for qPCR design are discussed.

## Introduction

Quantitative polymerase chain reaction (qPCR) has proved to be an important technology to give high sensitivity and specificity for diagnosis since early day of Coronavirus disease 2019 (COVID-19) ^1^ □. In the initial days, clinical diagnosis was relied up on, in addition to, less specific Computer-Tomography (CT) scans ^2^ □. Coronaviruses, and Nidovirales order in general, have the largest RNA genomes of all RNA viruses and due to high fidelity RNA replication and transcription machineries, the number of new mutations occurring per replication is very low ^3^ □. According to Worldometers.info, more than 1.6 billion tests have been carried world-wide for 7.8 billion world population by end of February 2021 ^4^ □. Due to resource constraints, kits that detect only one to two genes have also come to use ^5^ □, and this may increase the chance of false negative results in case qPCR fails to detect one or two genes due to some reasons. Studies have reported a lack of sensitivity for the Real-Time Reverse Transcriptase (RT)-PCR test used to diagnose SARS-CoV-2 ^6,7^ □. Although test sensitivity could be lowered by errors in methodology, instrument, and diagnostic kit, a decrease in PCR efficacy due to mutation in the primer or probe binding sites is very hard to account for ^8^ □.

Mercatelli et al have analyzed over 48 thousand SARS-CoV-2 genomes, obtained from the GISAID database, deposited from the beginning to June 26, 2020 and found over 350 thousand mutations in the viral genomes ^9^ □. Effect of mutations on qPCR sensitivities has been exemplified in a previous influenza pandemic ^10^ □. The impact of mutations on PCR sensitivity carried at community or country-level depends upon two factors: first, the relative propensities of the target gene areas to undergo mutation, and second, on the prevalence of such mutated clades/strains in the population. The first scenario is described as unique mutations and the second by all or prevalent mutations in this article. We designed this study to identify genes which had lesser mutability and could be used as reliable target sites for PCR.

## Results and Discussion

Based on number of unique SNPs per nucleotide, we found that NSP10 was the most conserved region (Figure 1). 3’ UTR and 5’ UTR were the least conserved regions with a high tendency to undergo mutation than the others. Similar results were seen when the prevalence of total (non-unique or prevalent) SNPs were compared among different genomic regions with NSP10 being the most conserved and 5’ UTR being the least. In general, non-structural proteins were more conserved compared to structural ones (Table 1). During the multiplication process of the viruses, the whole plus-stranded genomic RNAs are manufactured from minus-strand templates by a single type of replication machinery ^3^□, and thus the non-structural genes (open reading frame 1ab) and the non-structural genes (in the 3’ end of the coronavirus genome) should have similar mutation rates. While there could be different factors involved in this phenomenon, one of them could be evolutionary pressure for structural proteins to evolve. Structural proteins are exposed to antibodies in the respiratory mucosa or blood during infection and transmission from one cell to another. The non-structural proteins help in the intracellular physiology, particularly related to replication and transcription, and thus are unexposed to antibody containing fluids. Thus the structural proteins need to evolve to evade antibody-based suppression ^11^□ of infection to new cells.

**Figure 1:**
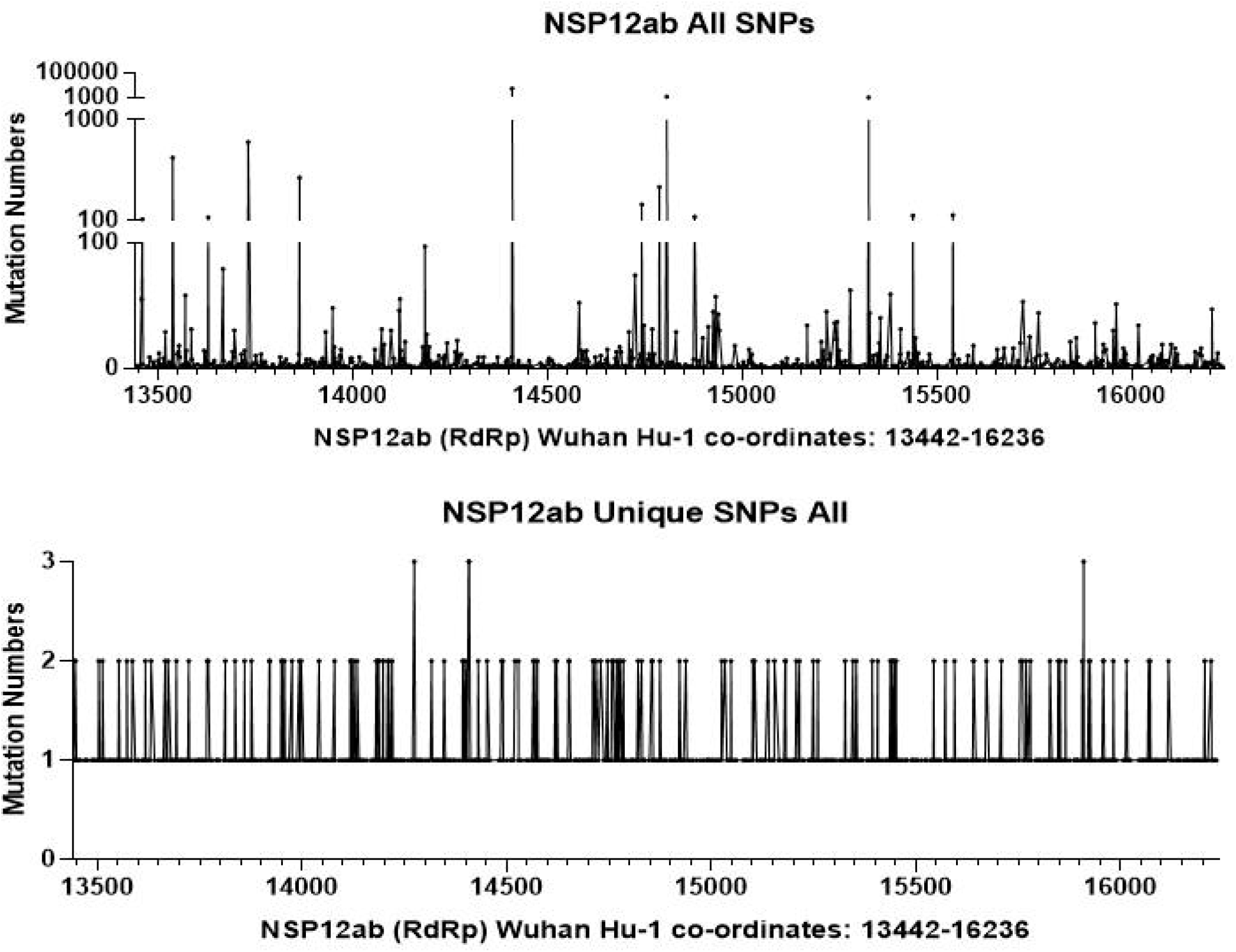
A.Unique SNP counts per nucleotide per 1 million genome. B. SNPs counts per Nucleotide per 1 Million Genome

**Table 1:** SNPs per nucleotide per million bases in different genes of SARS-CoV-2

| A |  |  |  | B |  |  |  |
| --- | --- | --- | --- | --- | --- | --- | --- |
| Genomic Region | Unique SNP Count | Region size | Unique SNPs/Nt /1Mil. Gen. | Genomic Regions | Total SNP Counts | Region size | Total SNPs/Nt/1 Mil. Gen. |
| NSP10 | 158 | 417 | 7.791 | NSP10 | 718 | 417 | 35.403 |
| NSP5 | 365 | 918 | 8.175 | NSP8 | 1238 | 594 | 42.853 |
| NSP8 | 239 | 594 | 8.273 | NSP16 | 2050 | 894 | 47.148 |
| NSP16 | 361 | 894 | 8.303 | NSP9 | 1179 | 339 | 71.51 |
| NSP12 (RdRp) | 1140 | 2795 | 8.386 | E | 837 | 228 | 75.482 |
| NSP13 | 756 | 1803 | 8.621 | ORF7b | 574 | 132 | 89.411 |
| NSP9 | 143 | 339 | 8.673 | NSP7 | 1112 | 249 | 91.824 |
| NSP14 | 706 | 1581 | 9.182 | ORF7a | 1664 | 366 | 93.481 |
| NSP4 | 679 | 1500 | 9.307 | ORF6 | 904 | 186 | 99.932 |
| NSP6 | 418 | 870 | 9.879 | NSP14 | 8351 | 1581 | 108.607 |
| NSP3 | 2818 | 5835 | 9.93 | NSP13 | 10084 | 1803 | 114.997 |
| NSP7 | 122 | 249 | 10.074 | NSP5 | 5176 | 918 | 115.932 |
| S | 1897 | 3811 | 10.235 | NSP1 | 3056 | 540 | 116.362 |
| M | 340 | 669 | 10.45 | NSP4 | 8696 | 1500 | 119.201 |
| NSP15 | 543 | 1038 | 10.756 | M | 4059 | 669 | 124.751 |
| ORF10 | 64 | 117 | 11.247 | ORF10 | 710 | 117 | 124.774 |
| E | 125 | 228 | 11.273 | NSP15 | 6470 | 1038 | 128.162 |
| ORF6 | 111 | 186 | 12.27 | NSP3 | 56466 | 5835 | 198.974 |
| NSP2 | 1157 | 1914 | 12.429 | NSP6 | 8697 | 870 | 205.542 |
| ORF7b | 85 | 132 | 13.24 | NSP2 | 22135 | 1914 | 237.787 |
| NSP1 | 351 | 540 | 13.365 | S | 51998 | 3811 | 280.543 |
| ORF8 | 256 | 366 | 14.382 | ORF8 | 6514 | 366 | 365.947 |
| N | 930 | 1260 | 15.176 | NSP12 (RdRp) | 50441 | 2795 | 371.068 |
| ORF7a | 277 | 366 | 15.561 | 3'UTR | 4738 | 229 | 425.413 |
| ORF3a | 631 | 828 | 15.669 | N | 27209 | 1260 | 444.01 |
| 5'UTR | 334 | 265 | 25.915 | ORF3a | 22717 | 828 | 564.12 |
| 3'UTR | 433 | 229 | 38.878 | 5'UTR | 37872 | 265 | 2938.485 |

Transcriptomic analysis of SARS-CoV-2 has repeatedly shown a higher prevalence of reads from the 3’ subgenomic RNAs in the infected cells ^12^ □. While this could be due to higher concentration of the sub-genomic RNAs located in the 3’ end, it was also reasoned that the methodological bias due to sequencing from the 3’ end of the genome might have impacted on the viral sequence reads ^12^ □. Sawicki et al have iterated that the cellular concentration of plus-stranded RNAs, which are synthesized by using minus-strand as a template, is 50-100 fold higher than the minus-stranded RNA for coronaviruses ^13^ □. The so called “nested sub-genomic structures” are formed during coronaviral transcription process. This means, whenever a gene, 3’ to the ORF genes, is transcribed for its translation, all other genes in 3’ direction to that particular gene are redundantly transcribed too. Considering that concentration of each of the plus-stranded translatable sub-genomic RNA units are 50-100 times more their corresponding minus-strands, this results in higher number of sequence reads for genes as we go towards the 3’ end of the genome causing the highest number of reads for N gene RNA followed by M, E and S ^12^ □. This could be the reason why the Ct values for structural genes N and E have better readings (lower Ct values) than that of the RdRp genes ^14,15^ □. Thus, the 3’ end sub-genomic RNAs for the structural proteins, which are present at higher concentration in the clinical samples, can be better regions for primer/probe design in terms of better analytical sensitivity.

Among the genes used by WHO collaborating laboratories, Corman et al (Charite, Gerrmany) used E gene and RdRp genes for the diagnosis of COVID-19 while other labs used N and/or ORF1 genes ^8^ □. Commercially, the S gene has also been used ^5^ □. “ORF1ab” has been frequently stated to be used for COVID-19 detection in the commercial PCR kits, but it is not clear if it is the RdRp gene (NSP12).

Different regions within specific genes may have different propensities to mutate depending on the exposure of different motifs to the antibody environment. We analyzed the mutation patterns (prevalent and unique) along 5’ to 3’ direction for major genes used in qPCR diagnosis (N, E, S and RdRp) (Figure 2). One can eyeball the mutation pattern along the length of the genes to select appropriate regions of each genes for primer/probe design. One should be careful to interpret that the shown mutation pattern in Figure 2 represent more than 45 thousand viral genomes and thus, any one clinical sample is highly unlikely to contain all the mutations. We would recommend low mutating structural genes to be used for diagnosis. Recently, the S-gene target failure in UK variant of concern strains tested by Applied Biosystems TaqPath RT-PCR COVID-19 kit was found to be due to 6-nucleotide deletion mutation in Spike gene region targeted by the kit ^16^□. Including only one gene such as this in qPCR diagnostic kit could have interpreted the clinical sample as negative. Thus kits with multiple targets are advisable. As the product sizes of individual genes in multiplex real-time PCR are maintained to be of equal or near equal and shorter lengths (100-200 bp) and of equivalent GC contents in addition to other factors ^17^ □, wise selection of regions with lower SNP loads should be carried to design reliable primers and probes.

**Figure 2:**
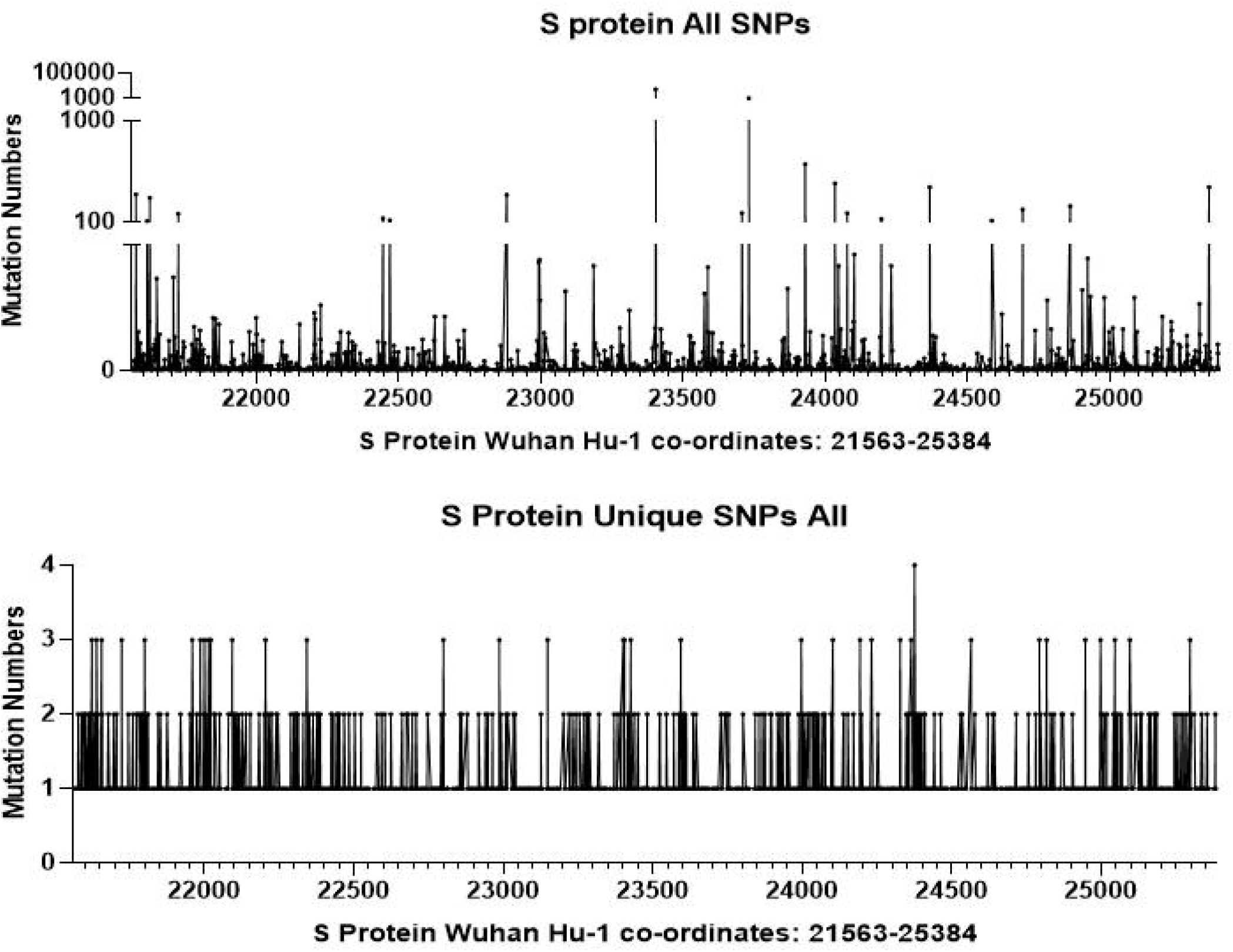
Visual representation of all and unique SNPs in N (A), E (B), S (C) and NSP12 (RdRp) (D) genes.

## Materials and Methods

Based on supplementary material provided by Mercatelli et al., we calculated the relative number of Single Nucleotide Polymorphisms (SNPs) per nucleotide in different genomic regions ^9^ □. While the data provided by Mercatelli et al. only included mutations until June of 2020, this analysis can help to understand the general trend of genes to mutate and give an idea about their relative mutability which is the objective of this study. Unique mutation events were determined, based on “refpos”, “refvar” and “qvar” columns in the excel, by removing the entries with duplicate SNP variants but retaining the SNP entries where the same nucleotide may have undergone different kind of mutations. To accurately quantify mutations in the 3’ UTR region, entries for sequences corresponding to 3’ UTR at or before nucleotide 29,674^th^, the last nucleotide for ORF10, were removed as the sequence only 3’ to ORF10 was included as true 3’ UTR. Intergenic SNPs, which were 3 in total, were also removed. We combined entries for NSP12a and NSP12b into NSP12 which corresponds to the RdRp gene. Number of mutations or SNPs, unique or all, per nucleotide per million genomes for each of the SARS-CoV-2 genes were calculated by dividing the number of SNPs in a given gene by product of nucleotide size of that given gene and number of genomes analyzed by Mercatelli et al. (48,635 genomes analyzed), and then multiplying 1 million. Figures were drawn in GraphPad Prism v5 or WPS excel. Gene co-ordinates of SARS-CoV-2 Wuhan-Hu-1 genome (NCBI accession ID NC_045512.2) was used as reference.

## Author Contribution

DRSJBR conceived the study. DRSJBR, NP and SD analyzed and wrote the manuscript. DRSJBR, NP and SD agreed on the final manuscript.

## Competing Interests

Authors declare they have no competing interests.

## Code availability

Mercatelli et al.^9^ □ used Bash/R codes to generate and annotate genome variants for genomes downloaded from GISAID.org. The Bash/R codes are available in the primary article (Mercatelli et al.).

## Data availability

Data are available in the supplementary material file 5 of Marcatelli et al^9^ □ (https://doi.org/10.3389/fmicb.2020.01800). Mercatelli et al downloaded data from www.GISAID.org. Excel files and Graphpad files used during analysis in this study are available on request.

---

Table 1: Number of all (A) and unique (B) SNPs in various genes of SARS-CoV-2 genome.

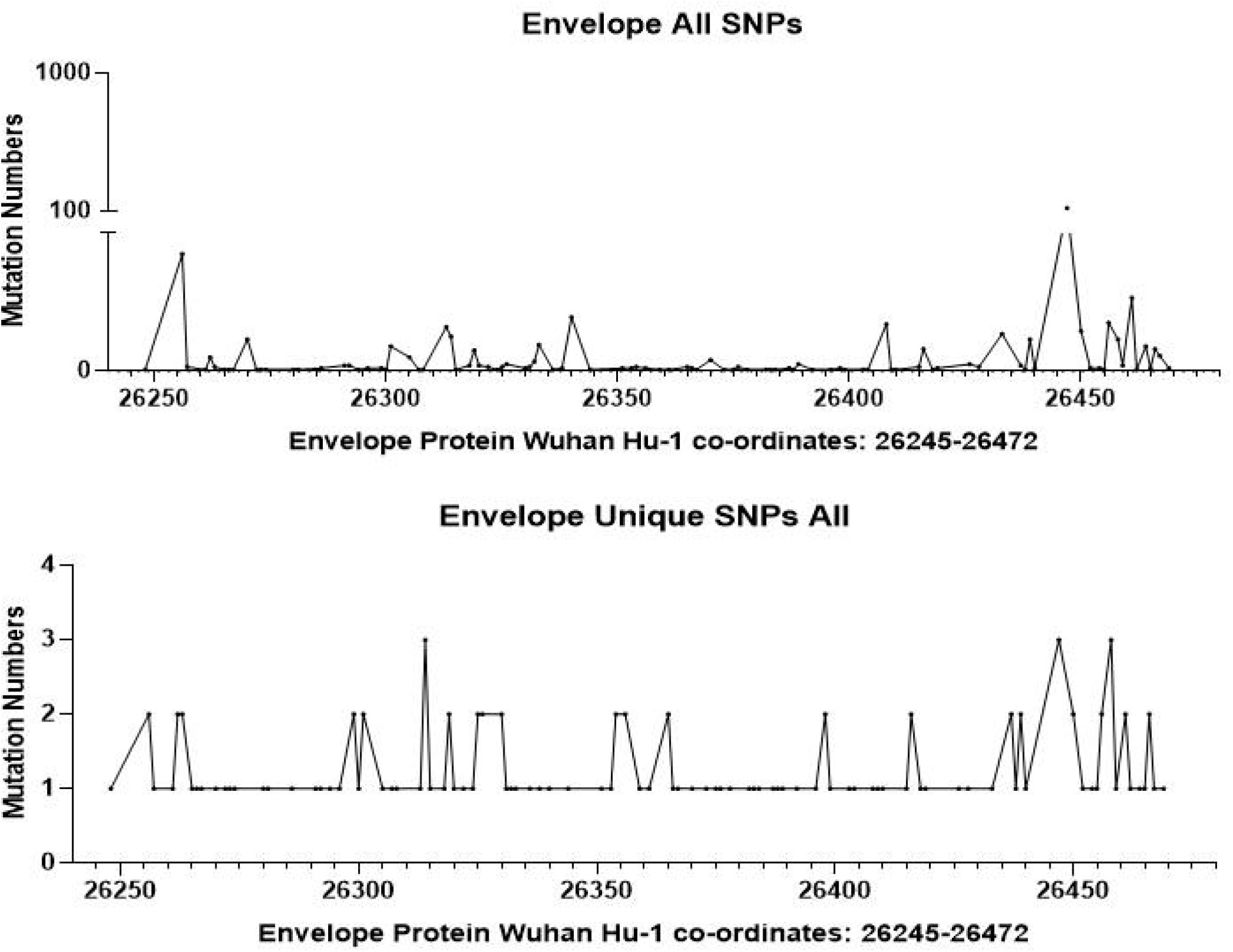

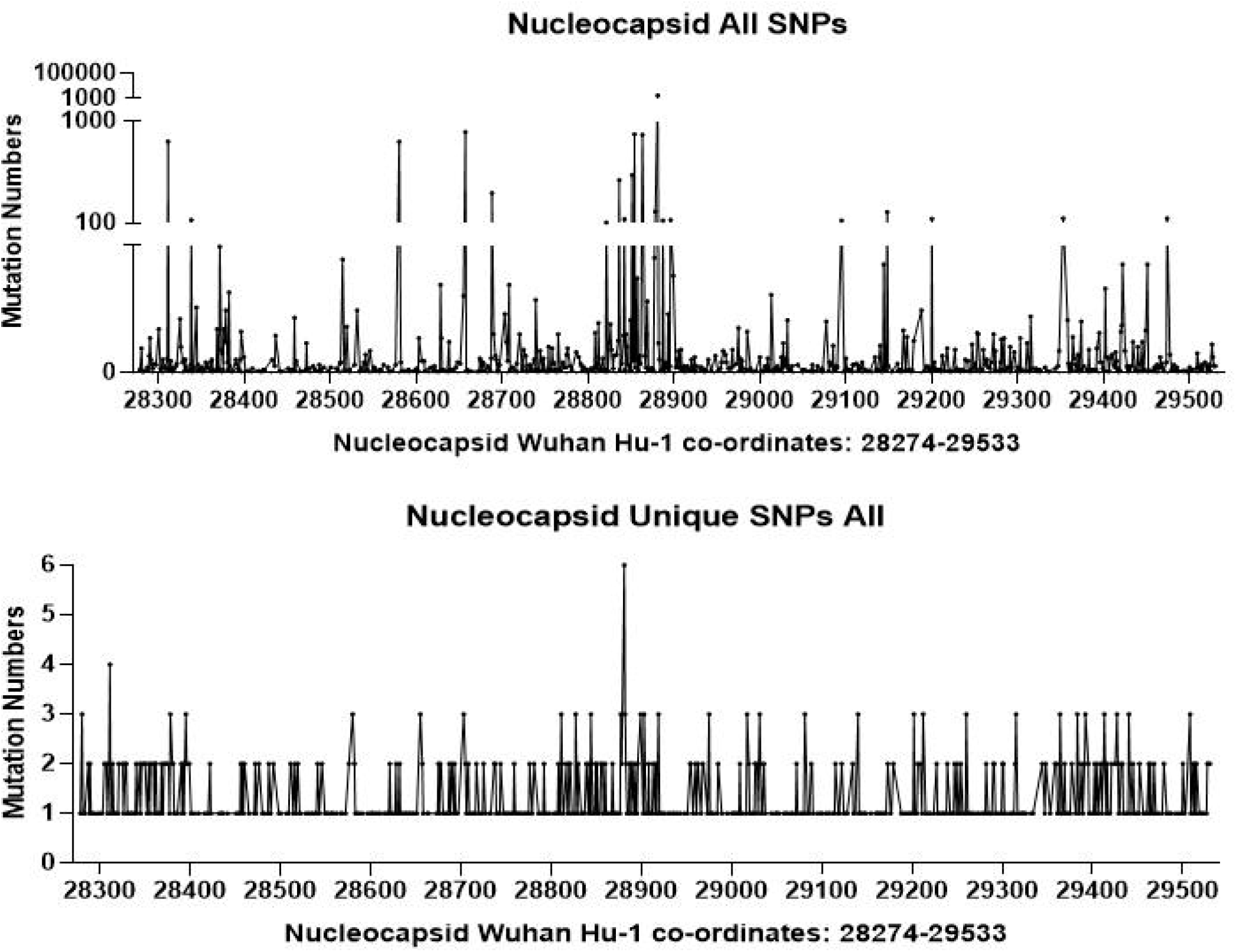

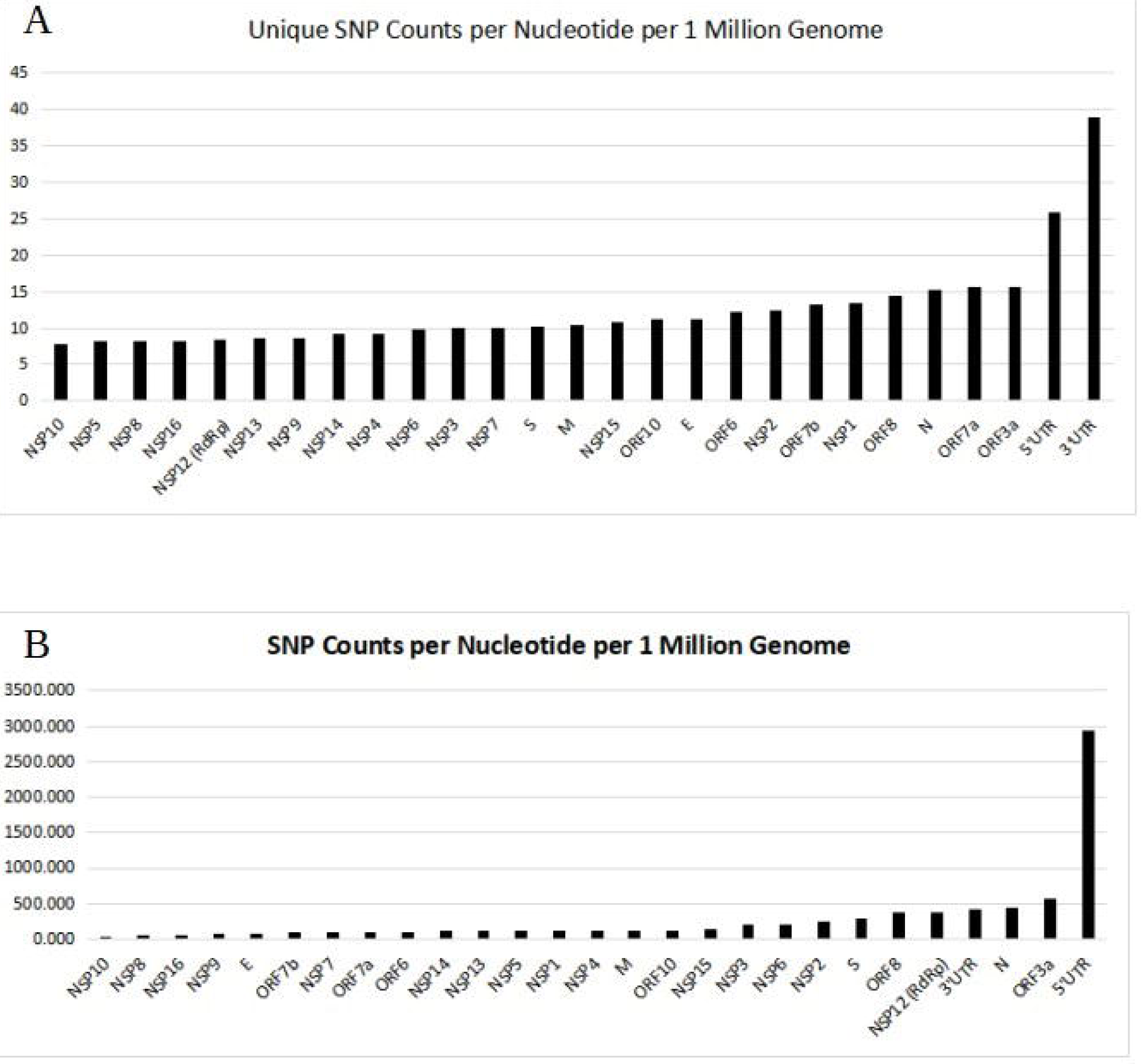

## Notes

### Competing Interest Statement

The authors have declared no competing interest.

## References

1. Corman, V. M. et al. Detection of 2019 novel coronavirus (2019-nCoV) by real-time RT-PCR. Eurosurveillance 25, (2020).

2. Ai, T. et al. Correlation of Chest CT and RT-PCR Testing in Coronavirus Disease 2019 (COVID-19) in China: A Report of 1014 Cases.

3. Sawicki, S. G. & Sawicki, D. L. Coronavirus transcription: A perspective. Current Topics in Microbiology and Immunology 287, 31–55 (2005).

4. Worldometers.info. Coronavirus Update (Live): 113,988,846 Cases and 2,529,393 Deaths from COVID-19 Virus Pandemic - Worldometer. (2021). Available at: https://www.worldometers.info/coronavirus/. (Accessed: 27th February 2021)

5. Iglói, Z. et al. Comparison of commercial realtime reverse transcription PCR assays for the detection of SARS-CoV-2. J. Clin. Virol. 129, (2020).

6. Alcoba-Florez, J. et al. Sensitivity of different RT-qPCR solutions for SARS-CoV-2 detection. Int. J. Infect. Dis. 99, 190–192 (2020).

7. Li, Y. et al. Stability issues of RT-PCR testing of SARS-CoV-2 for hospitalized patients clinically diagnosed with COVID-19. J. Med. Virol. 92, 903–908 (2020).

8. Vogels, C. B. F. et al. Analytical sensitivity and efficiency comparisons of SARS-CoV-2 RT–qPCR primer–probe sets. Nat. Microbiol. 5, 1299–1305 (2020).

9. Mercatelli, D. & Giorgi, F. M. Geographic and Genomic Distribution of SARS-CoV-2 Mutations. Front. Microbiol. 11, 1800 (2020).

10. Klungthong, C. et al. The impact of primer and probe-template mismatches on the sensitivity of pandemic influenza A/H1N1/2009 virus detection by real-time RT-PCR. J. Clin. Virol. 48, 91–95 (2010).

11. Kamp, C., Wilke, C. O., Adami, C. & Bornholdt, S. Viral evolution under the pressure of an adaptive immune system: Optimal mutation rates for viral escape. Complexity 8, 28–33 (2002).

12. Kim, D. et al. The Architecture of SARS-CoV-2 Transcriptome. Cell 181, 914-921.e10 (2020).

13. Sawicki, S. G., Sawicki, D. L. & Siddell, S. G. A Contemporary View of Coronavirus Transcription. J. Virol. 81, 20–29 (2007).

14. Nalla, A. K. et al. Comparative performance of SARS-CoV-2 detection assays using seven different primer-probe sets and one assay kit. J. Clin. Microbiol. 58, (2020).

15. Colton, H. et al. Improved sensitivity using a dual target, E and RdRp assay for the diagnosis of SARS-CoV-2 infection: Experience at a large NHS Foundation Trust in the UK. Journal of Infection 82, 159–198 (2021).

16. Bal, A. et al. Two-step strategy for the identification of SARS-CoV-2 variant of concern 202012/01 and other variants with spike deletion H69–V70, France, August to December 2020. Eurosurveillance 26, 2100008 (2021).

17. Bio-Rad. qPCR Assay Design and Optimization | LSR | Bio-Rad. (2021). Available at: https://www.bio-rad.com/en-np/applications-technologies/qpcr-assay-design-optimization?ID=LUSO7RIVK. (Accessed: 27th February 2021)

